# A non-invasive, fast on/off “Odourgenetic” Method to Manipulate Physiology

**DOI:** 10.1101/2022.11.25.517891

**Authors:** Yanqiong Wu, Shanchun Su, Xueqin Xu, Xincai Hao, Wei Lu, Xiaohui Li, Linhan Wang, Wei Tian, Yan Gao, Gang Cao, Changbin Ke

## Abstract

Manipulating molecular processes governing physiological functions has significant potential for clinical therapeutics and is an important approach to elucidate the cellular basis of physiological functions. Here, we designed a Odourgenetic system co-expressed *Drosophila* odorant receptor system (DORs) consisting of OR35a and OR83b, which were exclusively activated by their odor ligand, 2-pentanone. Applying 2-pentanone to DOR-expressing cells or tissues induced calcium influx and membrane depolarization. By inhalation of 2-pentanone, we successfully applied DORs to manipulate behaviour, control insulin secretion and regulate blood glucose and manipulate muscle contraction and associated limb movement. Because 2-pentanone rapidly enters the blood upon inhalation and leaves the body by exhalation, this odorant can be used with DORs to manipulate cellular function, and the manipulation can be terminated at any time. Such feature approach significantly improves the safety and controllability of DORs used in the clinic. Thus, the present study developed a non-invasive, controllable, fast on/off method to manipulate cellular activity and behaviour on a time scale of minutes.

---

Developing a rapid, controllable method for manipulating physiological functions has significant potential for clinical therapeutics and basic research. Optogenetics has provided optical control of neuronal activity at the millisecond time scale^1–5^. However, this approach requires direct optical access to brain tissue, which is difficult because blue light does not readily penetrate whole organisms; light must be delivered using costly specialized equipment such as custom blue light sources with fibre optics or two-photon illumination systems^6^. Additionally, given the need for invasive equipment implantation and sufficient power^7^, it is difficult to apply optogenetics for disease treatment in real clinical practice.

Chemogenetic designer receptors exclusively activated by designer drugs (DREADDs)^8^ are a powerful approach for remote and transient manipulation of cellular activity with no need for specialized equipment^9,10^. A recent study showed that metabolically derived clozapine arising from systemic clozapine N-oxide (CNO) administration is indeed the *in vivo* actuator of DREADDs^9^. Clozapine binds with high affinity to many receptors and has side effects such as behavioural inhibition and potentially fatal agranulocytosis^11^. Thus, the use of clozapine as a DREADD actuator in humans may result in undesirable side effects^9^. Converted clozapine reaches its maximal concentration at 2–3 hours after CNO treatment^12^, indicating that the effects of CNO on cellular activity are most likely to occur at this time point. These dynamic pharmacological profiles of CNO *in vivo* result in a long and uncontrollable process due to irreversible drug effects and metabolism, which limits the potential for emergency clinical applications such as seizure control^10^.

*Drosophila melanogaster* odorant receptor 35a (OR35a) belongs to the seven-transmembrane G-protein-coupled receptor superfamily and is a typical ligand-gated ion channel^13^; this channel forms a complex with the odorant-binding subunit odorant receptor 83b (OR83b)^14–16^. 2-Pentanone is the natural ligand of OR35a. In the present study, we co-expressed OR35a and OR83b in rodent tissues by viral transduction. We demonstrated that *Drosophila* odorant receptors (DORs) were activated by inhalation of 2-pentanone and effectively manipulated physiological processes and rodent behaviour on a time scale of minutes, indicating excellent controllability of DORs in practice. Here, we provide an easy-to-use, noninvasive, and spatiotemporally controllable approach to manipulate physiological processes. Because of the safety, availability, and cost-effectiveness of 2-pentanone, this “odourgenetic” approach has great potential for clinical therapeutics.

## 2-Pentanone induces calcium influx in DOR-expressing cells

First, we describe our scheme for DOR cloning, design and activation by 2-pentanone and the process through which physiological functions are manipulated by this system (Fig. 1a and b). To verify whether 2-pentanone bound to and opened these DOR channels on mammalian cells, OR35a and OR83b were expressed in a nonspecific manner in both *in vitro* and *in vivo* systems using a plasmid with a Ubc promoter and an mCherry reporter (Extended Data Fig. 2a). A GCaMP-expressing plasmid and a DOR-expressing plasmid were co-transfected into Neuro-2a cells (Fig. 1c and d) and HEK293T cells (Extended Data Fig. 2b). Calcium influx imaging experiments indicated that bath application of 2-pentanone elicited robust calcium influx in DOR-expressing Neuro-2a cells (Fig. 1c-f) and HEK293T cells (Extended Data Fig. 2b-f). A patch-clamp experiment using Neuro-2a cells showed that bath application of 2-pentanone induced depolarization of the membrane potential, indicating inward rectification by DORs (Fig. 1g and h). These results indicated that DORs modulated intracellular calcium levels by inward rectification and might therefore be useful for manipulating calcium-dependent cellular processes.

**Fig. 1.**
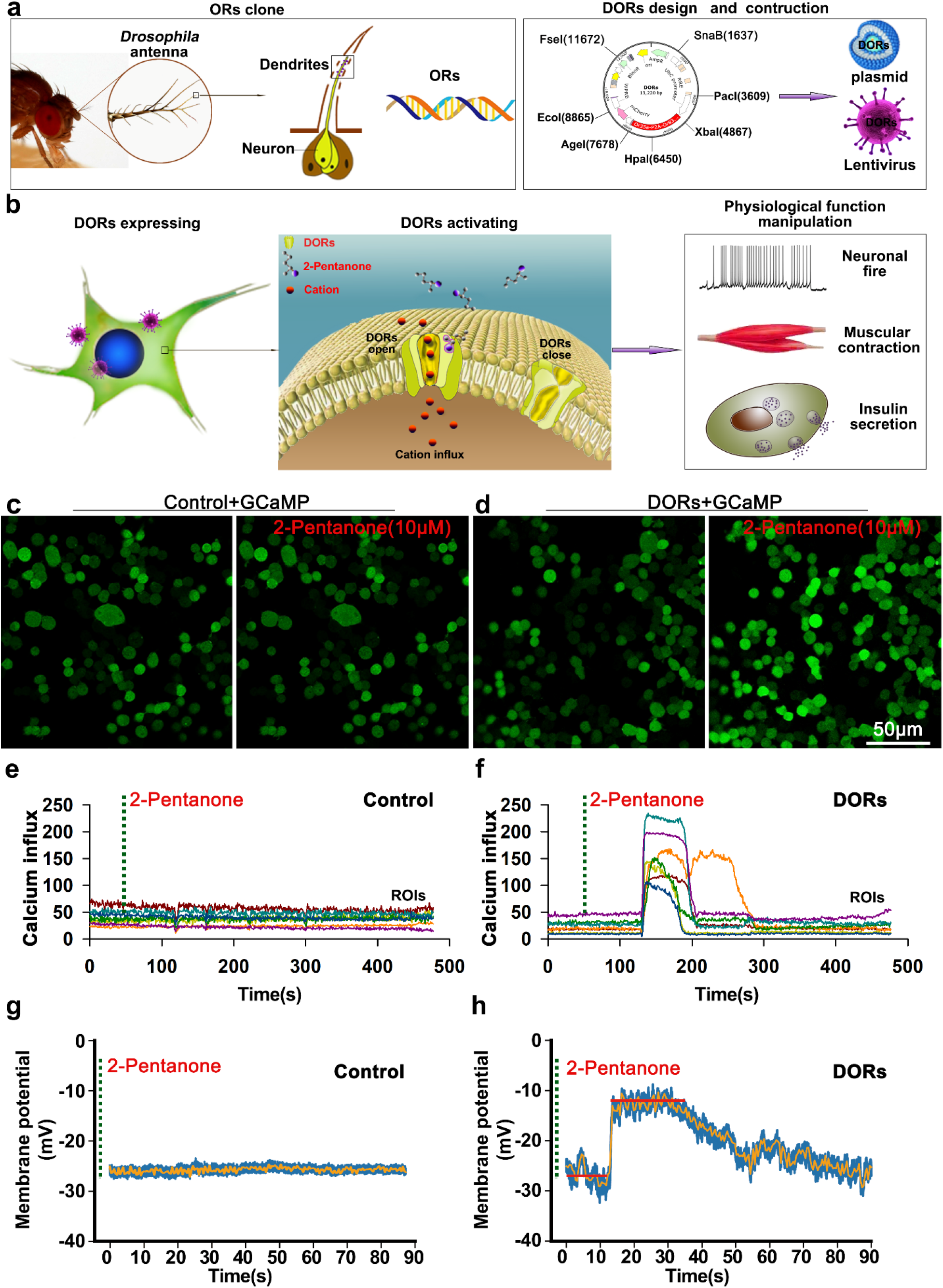
*Drosophila-derived* DORs control cellular activity. (**a, b**) Schematic drawing showing the principle of DOR design and their activation by 2-pentanone to manipulate physiological functions. (**c, d**) Two frames (before and after application of 2-pentanone) of time-lapse calcium influx images of GCaMP co-expressed with DORs or a control in Neuro-2a cells. The fluorescence responses showed that 10 μM 2-pentanone evoked significant calcium influx in DOR-expressing cells. (**e, f**) Time course of fluorescence responses from regions of interest (ROIs), showing a robust calcium influx in DOR-expressing cells in response to 10 μM 2-pentanone. (**g, h**) Changes in the membrane potential change of current-clamped Neuro-2a cells; bath application of 10 μM 2-pentanone significantly depolarized the membrane potential of DOR-expressing cells (mean change =16.38±7.79 mV, n=25 cells).

## DOR activation elicits spikes in DOR-expressing neurons

To investigate whether DORs manipulated neuronal activity, lentiviruses (LVs) encoding DORs and adeno-associated viruses (AAVs) encoding GCaMP were injected to the S1 cortex of C57 mice. Because the length of the DOR sequence is approximately 3.5 kb, a lentivirus vector was chosen for DOR expression in the present study. Neuronal spikes were recorded under current-clamp conditions in cultured neurons and acute brain slices. The fluorescence response to calcium influx was examined by confocal microscopy in acute brain slices. The results showed that 2-pentanone induced robust neuronal spikes in DOR-expressing neurons in cell culture and acute brain slices (Fig. 2a-d). Bath application of 2-pentanone elicited a continuous fluorescence response in DOR-expressing neurons of acute brain slices (Fig. 2e-n and Supplementary Video 1). These results indicated that DORs enabled 2-pentanone-driven manipulation of neuronal activity *in vitro* and *in vivo*.

**Fig. 2.**
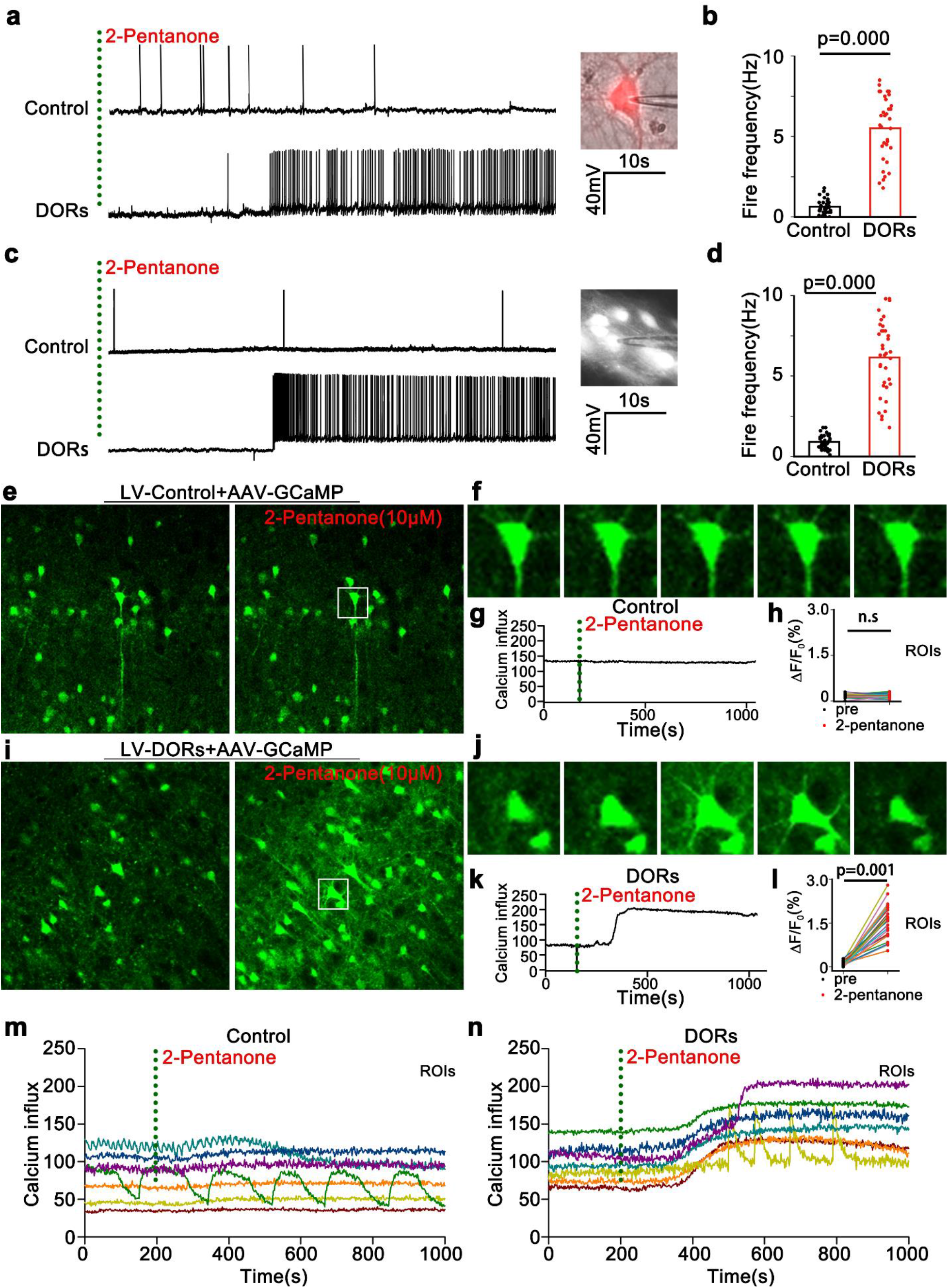
DORs enable 2-pentanone-driven neuronal spiking. (**a, c**) Voltage traces showing spikes in single current-clamped neurons of cultured cortical neurons and brain slices. Compared with the control, bath application of 10 μM 2-pentanone evoked robust firing spikes of DOR-expressing neurons in brain slices and cultured cortical neurons. (**b, d**) Spikes in current-clamped neurons in brain slices and cultured cortical neurons in response to application of 2-pentanone (slice: Control, 1.1±0.6 Hz, DORs, 6.9±3.1 Hz, p=0.000, n=36 neurons from nine mice; cultured neurons: Control, 0.8±0.5 Hz, DORs, 6.1±3.6 Hz, p=0.000, n=34 neurons from six cultures, non-parametric Mann–Whitney rank-sum test, two-sided). Experiments were repeated independently more than five times with similar results. (**e, i**) Two frames (before and after application of 2-pentanone) of time-lapse calcium influx images of brain slices from C57 mice co-infected with AAV-GCaMP and LV-DOR or LV-control. Compared with the control, 10 μM 2-pentanone evoked significant calcium influx in DOR-expressing neurons. (**f, j**) Snapshots of fluorescence responses of control or DOR-expressing sample neurons (marked with white rectangles on left images) to 2-pentanone. (**g, k**) Time course of fluorescence responses to 2-pentanone in the above neurons. (**h, l**) Mean fluorescence intensity change (ΔF/F_0_) of DOR-expressing and control neurons in response to 2-pentanone (DORs: 1.56% ± 0.18%; control: 0.18% ± 0.02%, p=0.001, n = 25 neurons from eight mice). (**m, n**) Time course of fluorescence responses of control and DOR-expressing neurons in ROIs during application of 2-pentanone. Compared with the control, DOR-expressing neurons showed enhanced fluorescence responses to 10 μM 2-pentanone.

## DORs enable fast on/off control of behaviour

For DOR expression under the control of the Cre-loxp system in the nervous system, an hSyn promoter following a loxp-stop-loxp sequence was inserted before the DOR sequence (Extended Data Fig. 2a). To determine whether DORs activated specific neurons, these receptors were expressed in GABAergic neurons in the S1 cortex of VGAT-Cre mice. Calcium influx imaging of brain slices was carried out to assess the fluorescence response of GABAergic neurons to 2-pentanone. The fluorescence response of brain slices showed that 2-pentanone activated the target DOR-expressing GABAergic neurons (Fig. 3a-c and Supplementary Video 4).

**Fig. 3.**
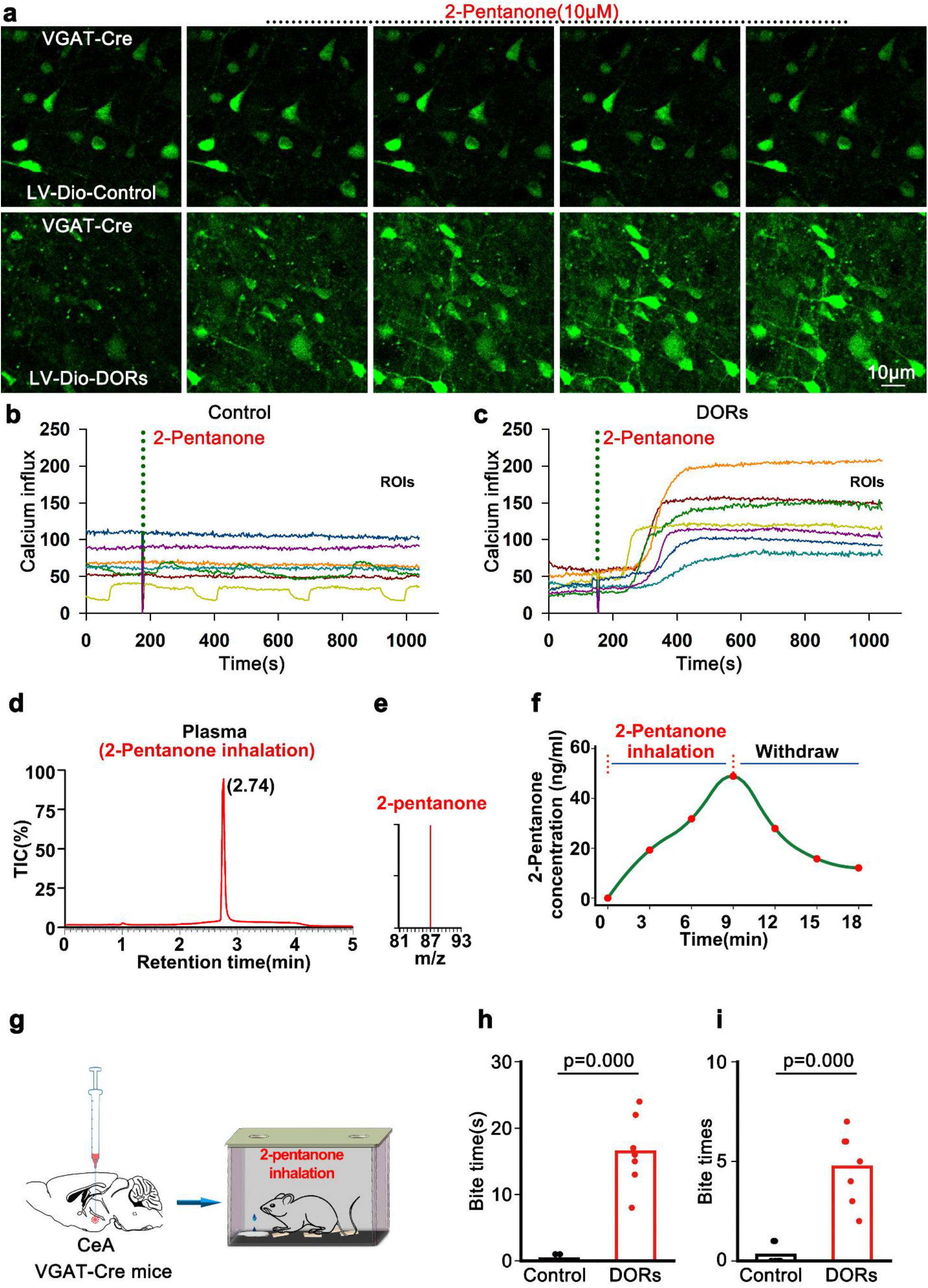
DORs activate GABAergic neurons in the CeA and control predatory-like behaviours. (**a**)Time-lapse calcium influx images of brain slices from VGAT-Cre mice co-infected with AAV-GCaMP and LV-hSyn-LSL-DORs or a control virus; bath application of 10 μM 2-pentanone evoked significant calcium influx in DOR-expressing neurons. (**b, c**) Time course of fluorescence responses of control and DOR-expressing neurons in regions of interest (ROIs) during application of 2-pentanone. Compared with the control, DOR-expressing neurons showed enhanced fluorescence responses to 2-pentanone. (**d, e**) The concentration of 2-pentanone in the blood of mice was examined by LC–MS after inhalation of 2-pentanone (2%, v/v) for 3 min. 2-Pentanone was identified by the retention time (approximately 2.74 min) and the mass charge ratio (m/z= 87). (**f**) Dynamic concentration of 2-pentanone in the blood of rats exposed to this compound. The 2-pentanone concentration showed a rapid time-dependent increase within a few minutes after inhalation, and the concentrations remained elevated until withdrawal of the stimulus. Upon withdrawal of 2-pentanone, concentration decreased to approximately 20% of the peak concentration within 10 minutes (48.88±10.63 ng/ml to 12.11±3.76 ng/ml, n=4 for each time point). (**g**) Schematic of virus injection into the CeA and inhalation of 2-pentanone in freely moving mice. (**h, i**) Values reflecting predatory-like behaviours evoked by 2-pentanone. Compared with the control, both the number of bites and the total time spent biting were increased in DOR-expressing mice (n= 6, non-parametric Mann–Whitney rank-sum test, two-sided).

Next, to determine whether 2-pentanone was delivered into the blood by inhalation and crossed the blood–brain barrier into the cerebrospinal fluid (CSF), we examined the 2-pentanone concentrations in the blood and CSF of mice exposed to this compound (2%, v/v) by LC–MS. Plasma containing 2-pentanone and pure plasma were used as positive and negative controls, respectively (Extended Data Fig. 1a-c). 2-Pentanone was detectable in both blood and CSF after a short period of inhalation, indicating that this odorant was transported into the blood and then to the CSF with this simple administration method (Fig. 3d, e and Extended Data Fig. 1d). Time-course examinations were carried out to explore the dynamic profile of the 2-pentanone concentration in the blood of mice and rats exposed to this compound. The 2-pentanone concentration in the blood of both rats and mice showed a time-dependent increase during inhalation of 2-pentanone and decreased rapidly after withdrawal of the odorant (Fig. 3f and Extended Data Fig. 1e). These results indicated that 2-pentanone had the appropriate profile to be a candidate manipulator of this DOR system.

The central nucleus of the amygdala (CeA) is a modular command system that exerts integrated control of predatory hunting in mice^17^, as we confirmed through an optogenetic experiment in the present study (Supplementary Video 5) to ascertain whether DORs can control predatory hunting behaviours by manipulating CeA neuronal activity. DORs were Cre-dependently expressed in GABAergic neurons in the CeA, and predatory-like bites induced by inhalation of 2-pentanone were observed. The behavioural experiment confirmed that 2-pentanone controlled rodent behaviours by exogenously expressing DORs in target neurons in a non-invasive, fast on/off manner (Fig. 3g-i and Supplementary Video 6).

## DORs enable fast on/off manipulation of physiological processes

To verify whether DORs reversibly manipulated physiological processes in mice, a lentiviral expressing DORs was injected into the pancreas, skeletal muscle and S1 cortex, respectively. 2-Pentanone was administered to the mice by inhalation; their blood insulin content was then examined using an ELISA, and their blood glucose was examined with a blood glucose meter. Furthermore, the contraction of virus-injected muscles activated by 2-pentanone was observed under a stereomicroscope, along with the associated limb movement. Neuronal activity was examined by *in vivo* calcium influx imaging. A few minutes of 2-pentanone inhalation resulted in a significant increase in the blood insulin concentration in pancreatic DOR-expressing mice and therefore lowered the blood glucose level (Fig. 4a-c). Inhalation of 2-pentanone elicited muscle contraction and limb movements within a few minutes, and the effect persisted until the odorant was withdrawn (Fig. 4d-f). Calcium influx imaging in the S1 cortex in vivo showed that 2-pentanone inhalation evoked robust continuous neuronal firing within a few minutes, which persisted until 2-pentanone withdrawal (Extended Data Fig. 3a-d and Supplementary Video 2). Repeated inhalation of 2-pentanone elicited another bout of neuronal firing. These results confirmed that DORs enabled reversible manipulation of several physiological processes *in vivo*.

**Fig. 4.**
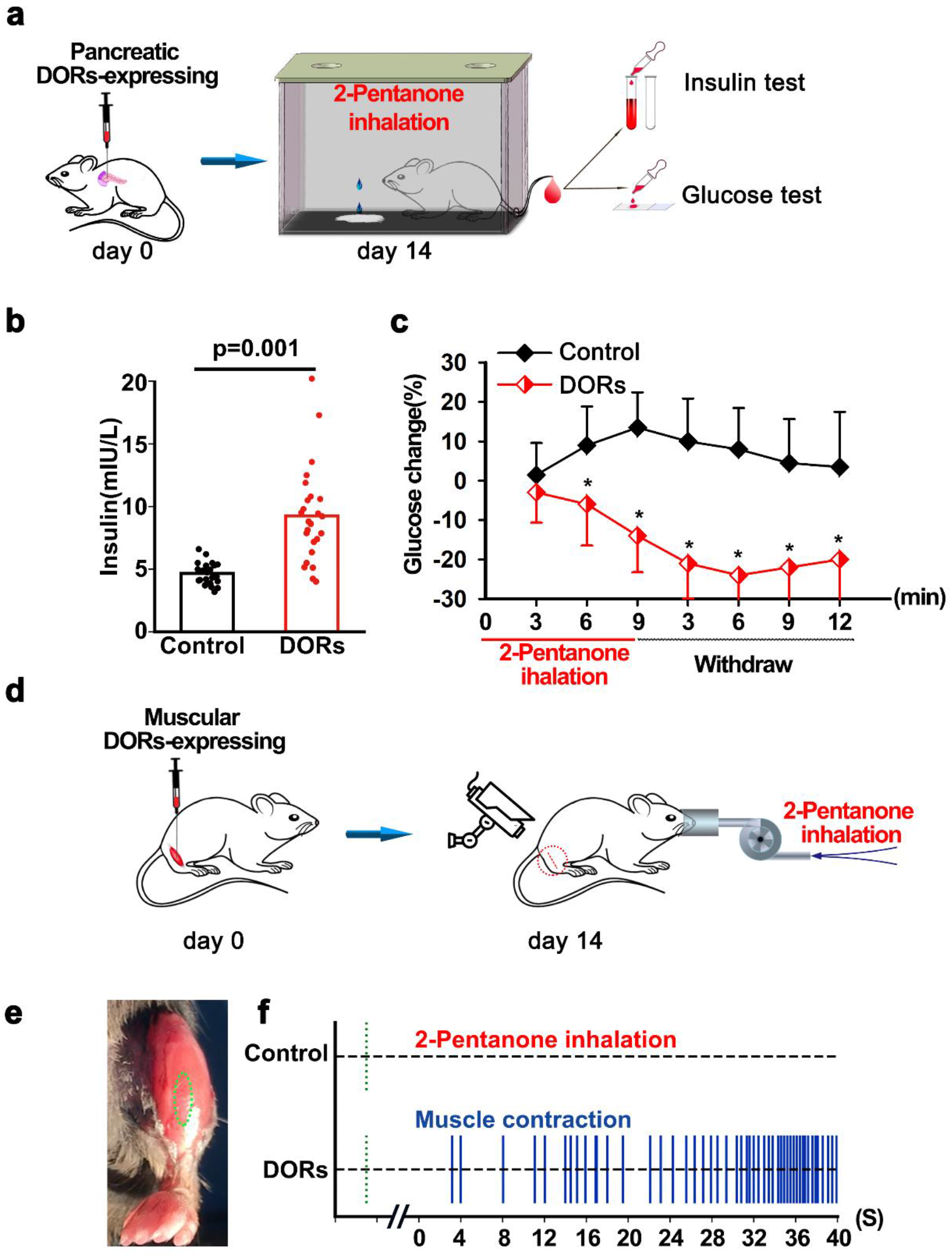
DORs manipulate physiological functions *in vivo*. (**a**) Schematic of virus injection into the pancreas and inhalation of 2-pentanone in freely moving mice. (**b**) Concentrations of insulin in the blood of mice after inhalation of 2-pentanone (control: 4.82±1.04 mIU/L; DORs: 9.32±3.34 mIU/L, p=0.001, n=27 mice, *t* test, two-sided) are shown. (**c**) Time course of the blood glucose change in mice subjected to 2-pentanone inhalation. Compared with the control, blood glucose began to decrease within a few minutes of 2-pentanone inhalation, and this change persisted for more than 10 minutes after withdrawal of 2-pentanone. Data are shown as the mean ± s.e.m. (n = 25, *p<0.05, repeated-measures ANOVA). (**d**) Schematic of virus injection into muscles and inhalation of 2-pentanone using a mask. (**e**) The image shows a virus-injected muscle marked by a green circle. (**f**) Vertical blue lines represent the muscle contraction evoked by 2-pentanone. 2-Pentanone elicited continuous contraction of muscles expressing DORs.

In the present study, DORs overcome many limitations of other methods, including the need for expensive specialized equipment; the difficulty of delivering light to widely distributed cell populations; the invasive procedures required to activate optogenetic systems in deep tissue; and the long, slow pharmacodynamics and irreversible metabolic processes of the designer drugs used in chemogenetics^6,9^. 2-Pentanone is a naturally produced phytochemical that is present in bananas^18^ and carrots^19^; this colourless liquid ketone has an acetone-like or intensely fruity odour. It is sometimes used in very small amounts as a food additive to impart flavour. 2-Pentanone is soluble in water and volatilizes rapidly to a gas at room temperature. Therefore, it was very easy to administer by inhalation to manipulate our DOR system. Furthermore, the compound is eliminated very quickly, mainly via exhalation, without a significant metabolic process. This profile indicates the good controllability of systemic 2-pentanone levels in the present DOR system, providing an easy-to-use tool that has potential for clinical applications in the treatment of various diseases, such as diabetes, Parkinson’s disease, and neocortical seizures^20^.

## Acknowledgements

This work was supported by the National Basic Research Program of China (81971060,32250018).

## Author contributions

Y.Q.W designed and carried out behaviour test, analyzed data and wrote the paper; S.C.S carried out electrophysiological and viral injection; X.Q.X carried out plasmid and viral construction and cellular expressing; X.C.H designed and carried out 2-pentanone examination; W.L carried out 2-pentanone examination; X.H.L carried out calcium imaging; L.H.W carried out blood insulin and glucose test; W.T carried out muscle contraction experiments. Y.G carried out cells and neuron culture; G.C designed the study and analyzed data; C.B.K designed the study, analyzed data and wrote the paper.

